# Advance Glycation End-products accelerate amyloid deposits in adipocyte’s lipid droplets

**DOI:** 10.1101/2023.11.06.565836

**Authors:** Roza Izgilov, Nadav Kislev, Eman Omari, Dafna Benayahu

**Affiliations:** Department of Cell and Developmental Biology, Faculty of Medicine, Tel Aviv University, Tel Aviv 6997801, Israel

**Author notes:** **Corresponding author:** Prof. Dafna Benayahu Department of Cell and Developmental Biology, Faculty of Medicine, Tel Aviv University, Tel Aviv 6997801, Israel. **Author Contributions:** Conceptualization, I.R., O.E., and B.D.; Data acquisition and formal analysis, I.R., O.E., and K.N.; funding and supervision, B.D.; methodology, I.R., K.N. and O.E.; visualization, I.R., and K.N.; writing original draft, I.R. and B.D.; Review and editing, all co-authors. All authors have read and agreed to the published version of the manuscript. **Competing Interest Statement:** The authors declare no competing interest.

**Keywords:** Adipocyte, Lipid droplet, Amyloid, AGEs, ATGL

## Abstract

Adipose tissue dysfunction is central to insulin resistance, and the emergence of type 2 diabetes (T2D) is associated with elevated levels of carbonyl metabolites from glucose metabolism. In this study, using methylglyoxal (MGO) and glycolaldehyde (GAD) carbonyl metabolites, induced protein glycation leading to misfolding and β-sheet formation and generation of advanced glycation end products (AGEs). The formed AGEs compromise adipocytes activity.

Microscopic and spectroscopic assays were used to examine the impact of MGO and GAD on lipid droplet - associated proteins. The results provide information about how glycation leads to the appearance of amyloidogenic proteins formation that hinders metabolism and autophagy in adipocytes. We measured the beneficial effects of metformin, an anti-diabetic drug, on misfolded protein as assessed by thioflavin (ThT) spectroscopy and improved autophagy. In vitro findings were complemented by in vivo analysis of white adipose tissue (WAT), where lipid droplet-associated β-amyloid deposits were predominantly linked to adipose triglyceride lipase (ATGL), a lipid droplet protein. Bioinformatics, imaging, and biochemical methods affirm ATGL’s role in β-sheet secondary structure creation. Our results highlighted the pronounced presence of amyloidogenic proteins in adipocytes treated with carbonyl compounds, potentially reshaping our understanding of adipocyte pathology in the context of T2D. This in-depth exploration offers novel perspectives on related pathophysiology and underscores the potential of adipocytes as pivotal therapeutic targets, bridging T2D, amyloidosis, protein glycation, and adipocyte malfunction.

**Significance Statement:** The generation of advanced glycation end products (AGEs) has a strong connection to diabetes severity . Adipose tissue is known to play a key role in the metabolic impairment and obesity associated with diabetes. We used the carbonyl compounds methylglyoxal (MGO) and glycolaldehyde (GAD) to create AGEs in adipocytes. The results of this study indicate that glycation not only affects cell metabolism and impairs adipocyte lipolysis, but also alters autophagy and increases protein amyloid deposits related to the membrane of lipid droplets. We identify the ATGL as a protein prone to β sheet alteration. consequently, ATGL emerges as a pivotal actor in lipid droplet metabolism and a prospective therapeutic target for T2D complications.

## Introduction

Adipose tissue plays a central role in obesity pathophysiology and this tissue dysfunction promotes insulin resistance and type 2 diabetes (T2D), a chronic metabolic disease characterized by high levels of blood glucose (hyperglycemia) (1, 2). Glucose metabolism generates a variety of carbonyl metabolites, including methylglyoxal (MGO) and glycolaldehyde (GAD). Methylglyoxal is also a byproduct of glycolysis which is impaired in T2D, results in its elevated levels (3, 4). Chronic hyperglycemia accelerates this process and exposes the cells to high levels of carbonyl metabolites, which react with lipids and proteins to generate advanced glycation end products (AGEs) through the Maillard reaction (5, 6). Protein glycation leads to alterations in protein structure and thereby promotes protein misfolding, β-amyloid formation and creates protein aggregation. Notably, such modifications occur in various proteins; for example, insulin and amyloid-β proteins, which have been associated with dysfunction due to protein alterations (7, 8).

Amyloidogenic proteins have the potential to create β-sheet structures when multiple β-sheets form fibrils, and the resultant unstructured protein assembles into insoluble fibers that accumulate and induce cell toxicity (9, 10). These cytotoxic amyloid aggregates are associated with a range of pathologies that are now recognized to extend beyond neurodegenerative diseases. Currently, around 50 proteins/peptides known to have the potential to deposit amyloid, including the T2D associated polypeptide amylin (IAPP) (11–13). In this context, we recently demonstrated alterations in the protein secondary structure and generation of amyloids after incubation of serum albumin with MGO and GAD (14).

The glycation byproducts promote protein structural modification and interfere with normal protein activity, leading to aberrations in the associated signaling cascade (8, 15, 16). Physiologically, such proteins should be recognized and eliminated by the autophagy system, which employs auto-phagosome formation and activation of the LC3 protein pathway. This highly regulated process, termed autophagy, is responsible for the degradation and recycling of cellular components and plays a critical role in maintaining cellular homeostasis. The process encompasses the clearance of misfolded or amyloid proteins, such as β-sheet aggregates, thereby preventing the accumulation of harmful entities that could promote pathologies such as in neurodegenerative disorders, Alzheimer’s and Parkinson’s diseases (17–21).

Chronic hyperglycemia in Type 2 Diabetes (T2D) has also been recognized to lead to insulin insensitivity by damaging pancreatic β-cells (22–24). In addition, an alteration in insulin signaling affects adipose tissue function leading to a vicious cycle of high glucose levels and consequently promotes hyper glycation and the formation of AGEs. The connection between AGEs formation and the deposition of amyloidogenic fibrils in adipocytes may be important in understanding the patho-physiology of adipose tissue. The glycation creates irreversible misfolded protein structures and the formation of aggregates as we recently have shown for serum albumin (14), and in an in vitro cell system under glycated conditions (25).

Lipid droplets are essential adipocyte organelles that store and release lipids to meet the energy demands of the organism. The role of lipid droplets in adipose tissue metabolism is well-established, but recent studies have shown that lipid droplets are also associated with a variety of proteins that play critical roles in lipid metabolism, adipogenesis, and metabolic homeostasis (26–29). The lipid droplet-associated proteins regulate various aspects of adipocyte biology, such as lipolysis, lipid uptake, and fatty acid oxidation, and are therefore critical for the regulation of adipose tissue metabolism. In addition, these proteins have been shown to interact with other cellular organelles, such as the endoplasmic reticulum and mitochondria, to coordinate lipid metabolism and energy homeostasis. Interestingly, alterations in lipid droplet function and size have been implicated in the development of adipocytes’ metabolic dysfunction. In a previous study we demonstrated that AGEs affect the level of adipogenesis (25), and here we use an advanced quantification approach to further examine the glycation effect on lipolysis and lipid droplet size (30).

Despite the strong association between T2D and amyloidosis, and although adipose tissue is a central player in insulin resistance and diabetes pathology, the formation of amyloidogenic proteins in adipose tissue has not yet been investigated. The current study was therefore designed to examine the effects of carbonyl compounds, glycation, and accumulation of amyloid structured proteins on the function of lipid droplets in adipocytes. The results provide new perspectives on amyloidogenic proteins and the conditions that promote such formation and affect the adipocytes’ function. We provide new information about the amyloid proteins that are associated with lipid droplets (LDs) and describe how hyper glycation impairs adipocyte function. We suggested that accumulation of misfolded proteins is a consequence of poor elimination by impaired autophagy as evidenced by MGO and GAD treated adipocytes and demonstrated the beneficial effect of the anti-diabetic drug metformin on this process. Among lipid droplet-associated proteins, we have identified ATGL as the primary protein that tends to promote the formation of amyloid deposits in cultured adipocytes as well as in visceral adipose tissue.

## Results

Pre-adipocytes have a flat elongated fibroblast-like shape (Fig. 1A left) that transitions morphologically to produce round mature adipocytes (Fig. 1A right) that contain lipid droplets (LDs). The accumulation of LDs is mediated by glucose uptake via the GLUT4 transporter, whose expression can be followed by assaying glucose uptake with 2NDBG. The expression of GLUT4 is higher on the adipocyte plasma membrane and peri-nuclear storage as reflected by a stronger fluorescent signal than that seen on the fibroblast (Fig. 1B). Glucose uptake using the 2NDBG followed by live-imaging shows a higher fluorescent signal in the adipocyte compared to fibroblast-like cells in the same culture (Fig. 1C, marked by a dashed line). The role of adipocytes in internalization of glucose motivated us to study the potential effect of hyper glycation due to the formation of AGEs.

**Figure 1:**
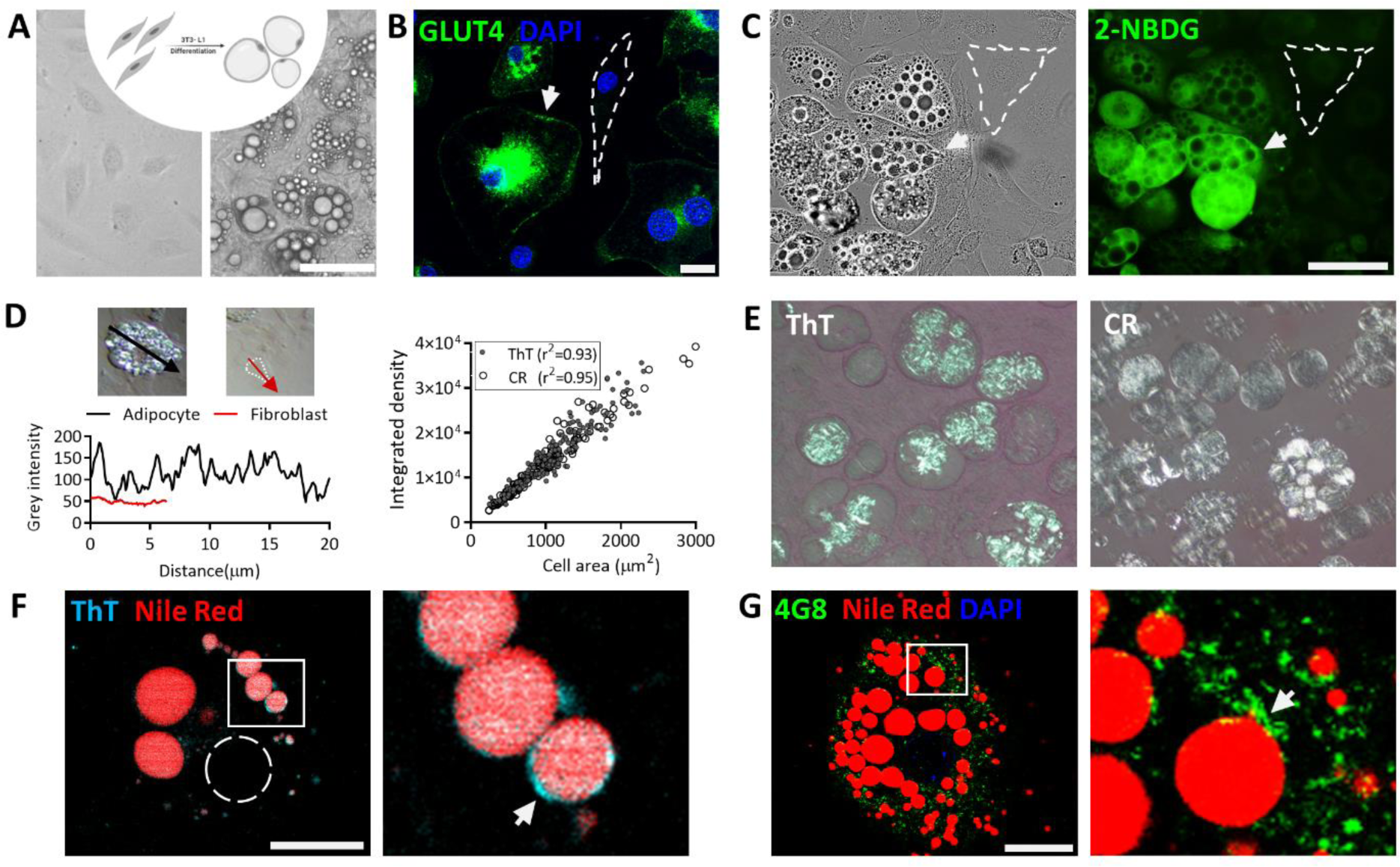
Adipocytes differentiation and amyloidogenic protein association with the lipid droplet. **A.** Schematic illustration of 3T3-L1 cell differentiation, pre-adipocytes (left) to adipocytes (right) and culture images (Magnification of x200, Scale bar= 125 µm). **B.** GLUT4 expression shown by IF staining, dashed line marks a fibroblast, an arrow points on membrane GLUT4 staining in an adipocyte. Magnification of X630. Scale bar= 20 µm. **C.** Phase contrast images of adipocytes (left) and 2-NBDG uptake (FITC, right) differentiate between pre-adipocytes (dashed line) and adipocytes (arrow). Scale bar= 100 µm **. D-E.** Polarized light microscopy images for Congo red (CR) and Thioflavin T (ThT) staining indicate amyloid formation (right panel). ThT staining Intensity as measured in fibroblasts (red) and adipocytes (black), the staining is quantified as a function of cell area (Simple linear regression). left panel; magnification x200; ThT, N = 175; CR, N = 145 cells. **F-G.** Confocal images of lipid droplet associated aggregates (arrows). Visualized by ThT probe (**F**) and 4G8 antibody IF (**G**). Lipid droplets stained by Nile-Red. Magnification of X630. Scale bar= 20 µm. Dashed line marks the cell nucleus.

We have previously reported that AGE formation increases the deposition of β rich amyloids and aggregates (14). Here, we monitored the effects on adipogenesis in 3T3-L1 cells treated with the carbonyl compounds MGO and GAD. Changes in protein conformation that reflect the development of amyloids can be monitored by staining adipocytes with Thioflavin (ThT) and Congo red (CR) and visualizing the resultant bright deposits with a polarized microscope (Fig 1D-E). This is the gold standard for measuring β amyloid formation. Notably, the adipocytes exhibit bright staining while the fibroblasts remain dark. The brightness is associated with the adipocyte lipid droplets, although not all LDs exhibit the same level of staining. The basal level for fibroblast-like cells was 50 IU, while the adipocyte levels were four-fold higher, predominantly located in the membrane of lipid droplets, and with a positive correlation between cell area and integrated density (Fig. 1D). These results were verified by co-staining of Nile-Red (NR) with ThT (Fig. 1F) and by co-staining by IF with the 4G8 antibody (green), that has a structure specific binding to the β sheet alteration in amyloidogenic proteins (Fig. 1G) (31).

The involvement of adipocytes in the internalization and metabolism of glucose led us to examine how the formation of AGEs can affect cellular activity and specifically the differentiation capacity of 3T3-L1 cells. As already discussed, we have previously reported that treating cell cultures with the carbonyl compounds MGO and GAD decreases the level of adipogenesis (LOA) (25). Here, we extended the study to changes in adipocyte morphology and specifically to effects on cellular area and LD size. Single-cell analysis enables to characterize the changes in LDs accumulation and cell morphology (Fig 2), did not detect any differences in cellular area. In contrast, incubation with MGO or GAD increased the LDs radius by 40%, with values of 4 µm ± 0.06 or 5.4 µm ± 0.09 after exposure to MGO or GAD respectively, compared to a mean radius of 2.9 µm ± 0.05 in untreated adipocytes (Fig 2A-C). The frequency distribution of LD size was also increased after treatment (Fig 2D).

**Figure 2:**
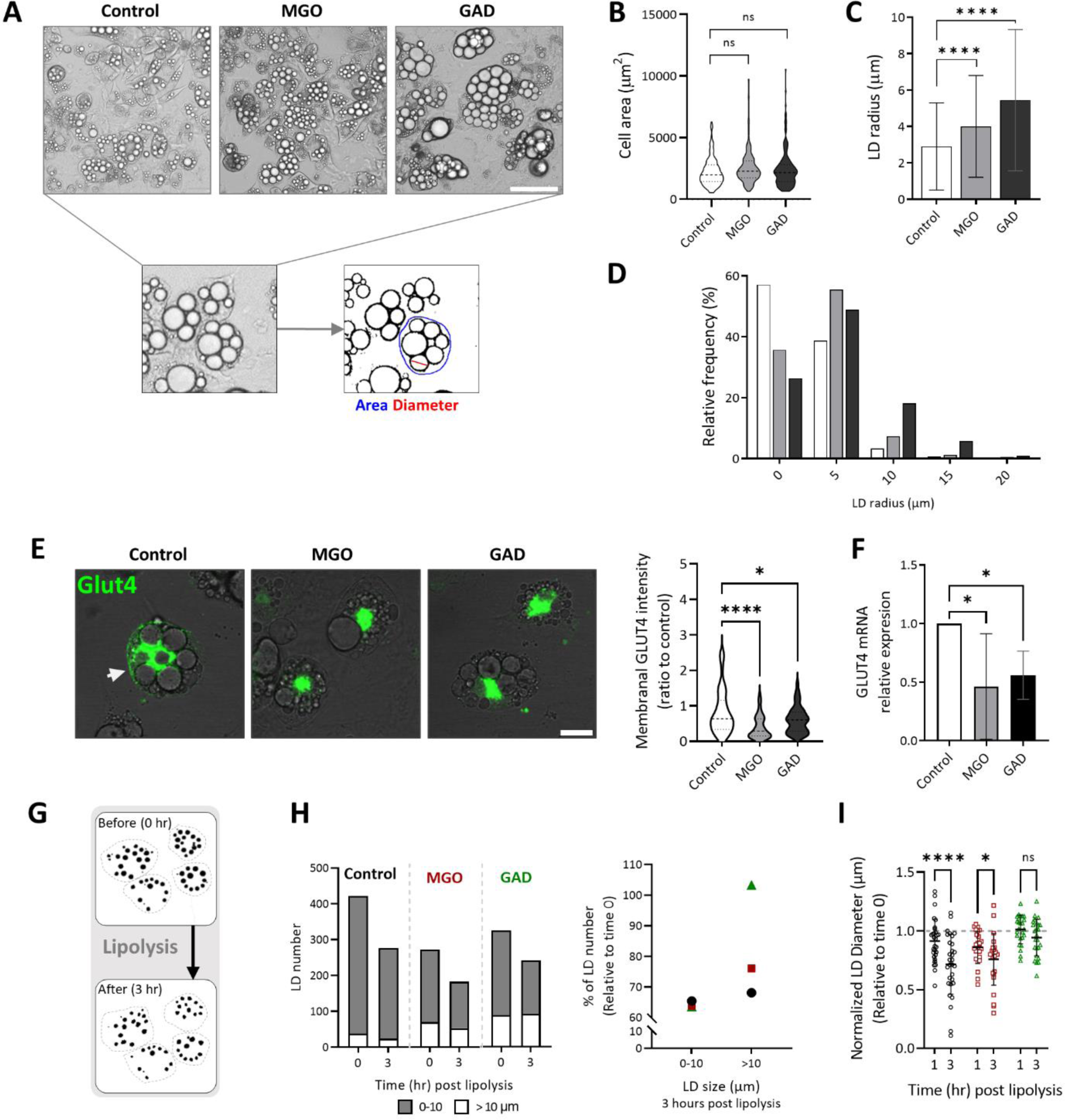
Carbonyl metabolites (MGO and GAD) influence adipocyte metabolism. **A.** Phase contrast images of adipogenesis for control, MGO and GAD treated cultures. Magnification x200; Scale bar= 125 µm. **B.** Measured cell area. **C.** Mean LD radius and size distribution (**D**) of untreated and treated adipocytes. (Control-white; N = 114, MGO-gray; N = 155, GAD-black; N = 184 cells). **E.** IF staining for GLUT4 membrane expression and quantification in MGO/GAD treated cultures (Magnification of X630. Scale bar = 20 µm, N = 50 cells/group). **F.** mRNA levels of GLUT4 in MGO and GAD treated adipocytes compared to controls (N = 4). (**B-F** One-way ANOVA). **G.** Illustration of changes in LD content at the single cell level during lipolysis induction (dashed line-cell membrane). **H.** Lipid droplet analysis during lipolysis measured by live imaging. Total LD number before and after 3 hours of lipolysis induction in MGO and GAD treated adipocytes. The LDs are divided into two groups by diameter size (left panel; White - large LD, > 10 μm; Gray - small LD, 0 < 10 μm). LD number change (% from the starting point) after 3 hours of lipolysis induction in small and large LDs (right panel; ●-control; ▪-MGO; ▴-GAD). **I.** LD diameter after 1 and 3 hours of lipolysis induction relative to the starting point. Two-way ANOVA. Data are presented as mean ± SD. (ns) p > 0.05; *p < 0.05; ****p ≤ 0.0001.

The GLUT4 transporter depends on insulin for glucose internalization into adipocytes, and transporter translocation to the cell membrane from cytoplasmic storage vesicles is a crucial step in the process. Fig 2E presents the results of staining cells for the presence of GLUT4. Quantifying the expression on cell membrane indicated that treatment with MGO and GAD reduces the GLUT4 expression on the adipocyte plasma membrane by a factor of two. This was accompanied by a decrease in GLUT4 mRNA level in the treated cells (Fig 2F). A reduction in GLUT4 transcription and cell membrane expression reflects the development of insulin signaling impairment. In addition to glucose uptake, an alteration in cell signaling also affects adipocyte lipid metabolism and storage by lipolysis. This was investigated by inducing lipolysis in treated adipocytes and observing the morphological changes over 3 hours by live imaging (Fig 2G-I). If the LDs are categorized as small or large by radius size (0-10 and >10 µm), the smaller LDs proved more sensitive to lipolysis induction than the large LDs regardless of treatment. However, 8% fewer large LDs reacted to lipolysis in MGO treated adipocytes, while GAD had no effect on the number of large LDs compared to control (30% less, Fig 2H). This profile is obvious from the change in LD diameter presented in Fig 2I, with a significant reduction in control cells compared to a moderate change in MGO treated cultures and no change in the cells treated with GAD. The proteins responsible for lipolysis are the LD-associated proteins that interact with the LD membrane, and this fact combined with the amyloid formation in Fig 1, led us to look for changes in amyloid formation in adipocytes treated with carbonyl compounds.

The lipid droplets were co-stained with ThT and with anti-plin-1, a lipid droplet membrane protein. ThT staining is related to LD size and is at higher level on the large LDs, although not all LDs are stained with ThT (Fig 3A-B). The heterogeneity in staining may be attributed to the level of amyloid formation during LDs maturation. Further, the effect of carbonyl metabolites on LDs were assessed at two time points: 7 and 14 days of treatment (Fig 3C-F), and we analyzed LDs diameter correlated with the presence of amyloid aggregation. Amyloids formation was detected as ThT^+^ by a polarized microscope. No changes in the level of amyloid positive LDs were detected in control cells over the course of the experiment. In contrast, treatment with MGO or GAD increased the levels of ThT^+^ LDs at both time points: MGO (from 66.6% to 78.2%) and GAD (from 65.7% to 92.0%, Fig 3C). Images of ThT^+^ stained LDs at day 14 are presented in Fig 3D-F, which presents the difference in ThT staining and the amyloid accumulation after treatment with MGO and GAD.

**Figure 3:**
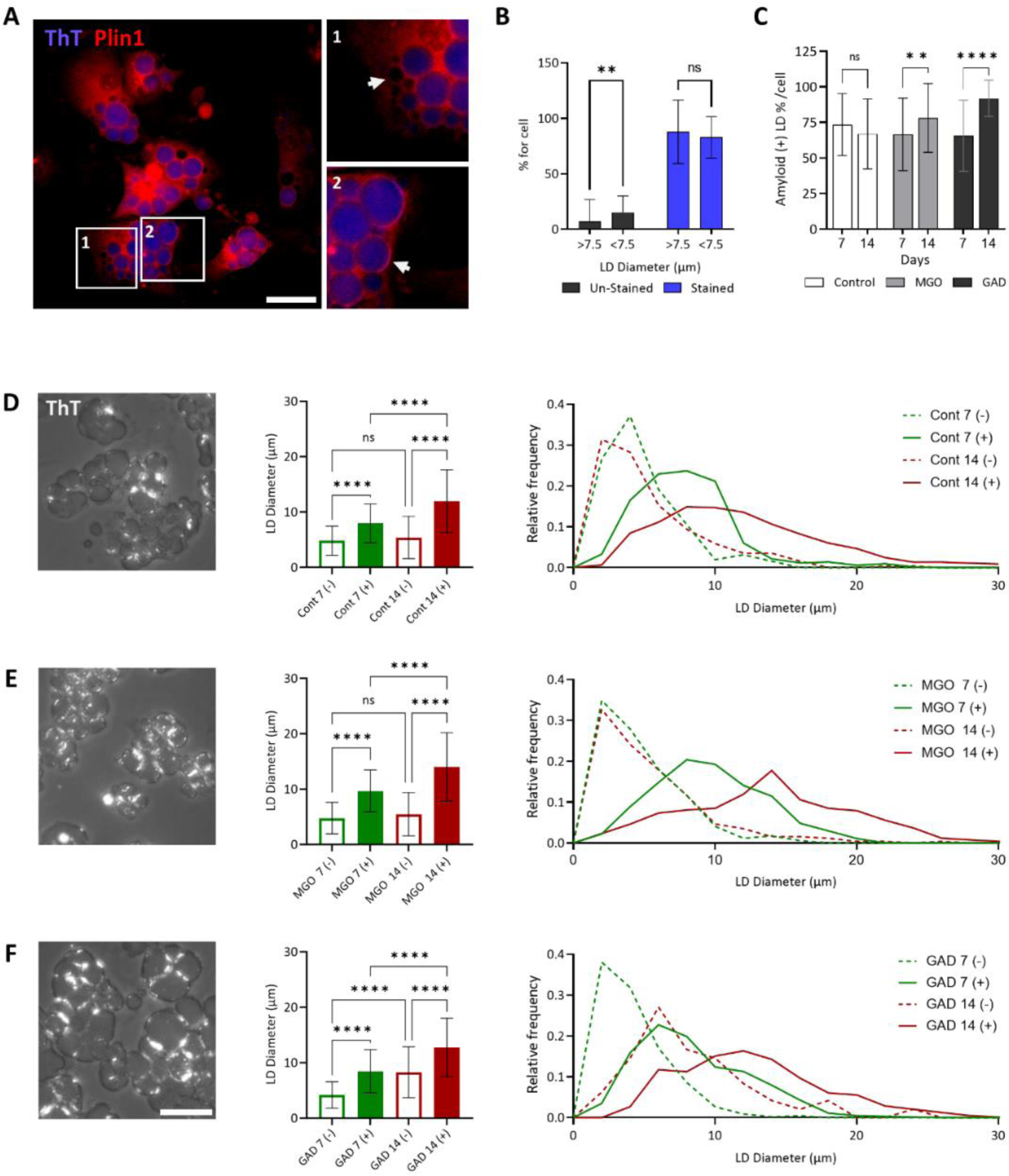
Amyloid-positive appearance on lipid droplets following exposure to carbonyl metabolites. **A.** Adipocytes stained for perilipin 1 (plin-1) and ThT: 1. Unstained LD (marked by an arrow); 2. Stained LDs with co-localization of ThT and IF for plin-1 in the LD membrane (marked by an arrow, magnification of x 200. Scale bar= 50 µm). **B.** Quantification of the percentage of ThT+ stained and unstained LDs (Diameter: large LD, > 7.5 μm; small LD, < 7.5 μm; N = 156 cells). **C.** Quantification of LD with ThT+ staining of amyloid (polarization images in D-F) after 7 and 14 days with MGO or GAD. Data are quantified as % of total LD count per-cell. Two-way ANOVA with Tukey’s post hoc test. **D-F.** Polarized images of ThT stained adipocytes after 14 days treatment (left), LD diameter quantification (middle panel) and distribution (right). The quantification of ThT+ and ThT-after 7 (green) and 14 (red) days of treatment - control (**D**); MGO (**E**); GAD (**F**). Scale bar= 50 µm. One-way ANOVA. Data are presented as mean ± SD. (ns) p > 0.05; **p ≤ 0.01; ****p ≤ 0.0001.

Similarly, the distribution of LD diameter at the two time points (Fig 3D-F) detected no difference in the distribution of ThT^-^ LDs in control cells over the course of the experiment, although the diameter of the ThT^+^ LDs increased after 14 days (from 7.9 µm to 11.9 µm). MGO treatment caused a significant shift only in the ThT^+^ LDs after 14 days (from 9.7 µm to 14.0 µm), and it became clear that the ThT^-^ LDs in both the control and MGO treated samples represent the smaller LDs (peak around 4-7 µm). In contrast, GAD treatment resulted in a size shift in both ThT^+^ (from 8.5 µm to 12.7 µm) and ThT^-^ (from 4.2 µm to 8.3 µm) LDs with time. These results support the LD radius differences and lipolysis results described above (Fig 2) and demonstrate a variation in the LD-amyloid aggregation caused by MGO or GAD.

The ThT^+^ staining between the treated groups indicates the β-sheet/amyloid accumulation associated with the LDs were further verified by RAMAN spectral analysis. Fig 4A show the images taken on RAMAN microscopy of untreated and MGO/GAD treated adipocytes. Further spectral analysis allows us to analyze the structural and chemical “fingerprint” of the proteins and can detect any alterations in chemical composition and molecular properties. Intracellular changes in the secondary structure of proteins caused by MGO/GAD treatment are reflected by the molecular profile in the measured spectrum. The results revealed that untreated or treated adipocytes have a similar peak at 1440 cm^-1^ which reached 0.99, indicating that all groups have the same lipid content (Fig 4B). However, significant differences in the chemical bonding and intramolecular bonds could be observed by analyzing the amide I peak at 1655 cm^-1^, which is associated with the secondary structures of proteins. The peak intensity was 1.3-fold higher in adipocytes treated with MGO or GAD than in untreated controls (Fig 4D). Similarly, the RAMAN signature at 1260 cm^-1^, which represents amide III and intramolecular β-sheet structures (Fig 4C), was 1.8-fold stronger in adipocytes treated with GAD compared to untreated adipocytes (Fig 4E). The results represent the differences in the secondary structures present in treated adipocytes.

**Figure 4:**
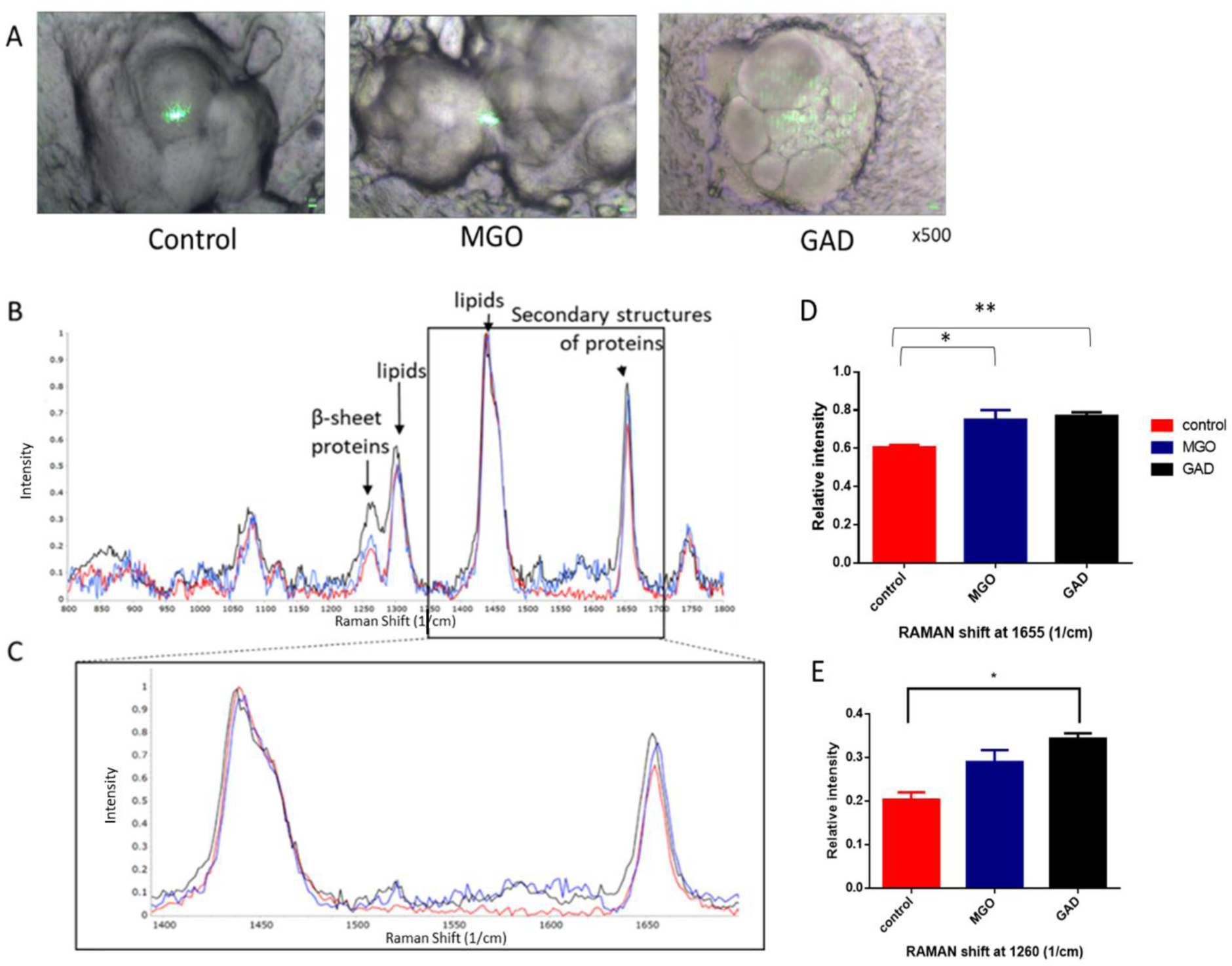
RAMAN spectra of carbonyl metabolites treated live adipocytes. **A.** RAMAN microscopy bright-field images of untreated and MGO/GAD treated adipocytes (Magnification of x500). **B.** RAMAN spectral profile comparison of untreated and treated adipocytes measured with a 532 nm excitation laser. Structure specific peak positions are labeled. **C.** Magnification of the spectral range 1400-1700 1/cm. **D.** Normalized intensity of RAMAN band at main peak at 1655 1/cm represents the secondary structures and the peak at 1260 1/cm (**E**) represents the β-sheet structure (N = 5-10 scanned areas). Control (red); MGO (blue); GAD (black). Data are presented as mean ± SD. One-way ANOVA, *p < 0.05; **p ≤ 0.01.

### Carbonyl compounds influence the intracellular degradation pathway

Conformational changes, such as the formation of β-sheets, can cause proteins to become amyloidogenic, which is known to alter protein function. Such a protein becomes cytotoxic and must be eliminated/degraded to maintain proper cellular activity. The removal of misfolded proteins is accomplished by the autophagy pathway. High resolution TEM images of the auto-phagosome in MGO/GAD treated cells (Fig 5A) reveal that the double membrane structure of the auto-phagosome (white arrow) that surrounds the cargo is dedicate to lysosomal degradation. Vesicles in cells grown with GAD appear larger and are characterized by complex cargo content. To better understand these observations, cultures were treated for 6 days with 5 μM metformin (MET), which is an inducer of autophagy (Fig 5B).

**Figure 5:**
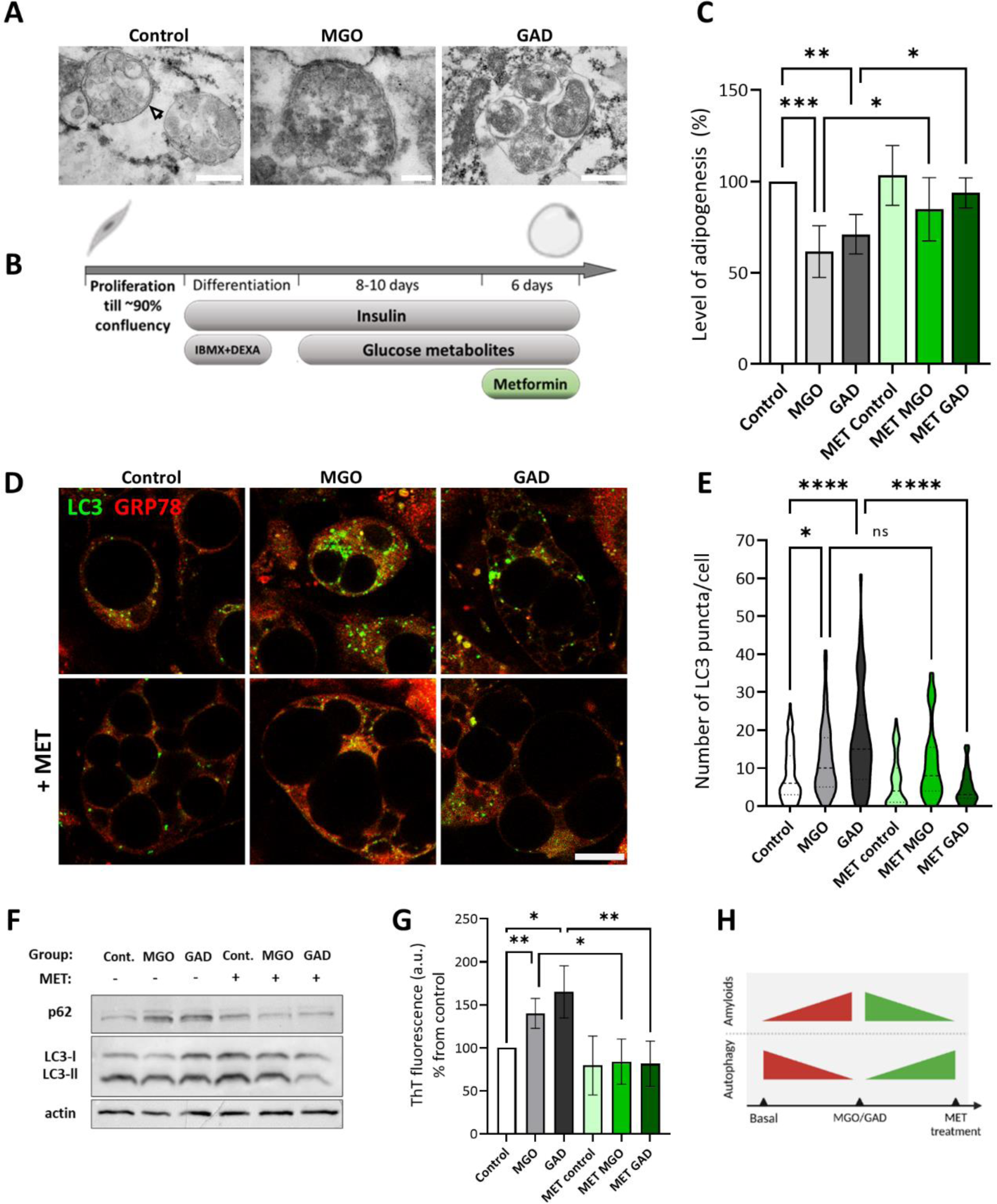
Metformin impact on autophagy in MGO and GAD treated adipocytes. **A.** TEM images of autophagy vesicles formed upon treatment with MGO and GAD. White arrow points to the autophagosome double membrane (Magnification and scale bar: control-X50k, 500 µm; MGO-X100k, 200 µm; GAD-X60k, 500 µm). **B.** Schematic illustration of experimental timeline with metformin (MET) treatment. **C**. LOA quantification of MGO and GAD treated adipocytes compared to controls (N = 6). Green coloring represents MET treated groups. **D.** Confocal images of autophagy puncta stained with anti-LC3 in MGO/GAD treated adipocyte cells with MET treatment (Magnification of X630, scale bar = 20 µm). **E.** Quantification of LC3 puncta number per cell in treated adipocytes (control, N = 89; MGO, N = 57; GAD, N = 68; MET, N = 96; MET+MGO, N = 59; MET+GAD, N = 85 cells). **F.** P62 and LC3 I/II protein expression levels in adipocytes treated with MGO, GAD, and MET. **G.** ThT fluorescent spectroscopy of MGO/GAD and MET treated adipocytes cell lysates. Data are presented as mean ± SD. One-way ANOVA with Tukey’s post hoc test, (ns) p > 0.05; * p < 0.05; **p < 0.01; ***p < 0.001; ****p < 0.0001. **H.** Schematic summary of MET effect on autophagy and amyloid levels in MGO and GAD treated adipocytes.

We have previously reported an altered LOA in cells treated with MGO/GAD (25). This was extended here to an examination of the effects of the carbonyl compound on lipolysis (Fig 2), the formation of amyloids by imaging ThT^+^ LDs (Fig 3), and by RAMAN spectroscopy (Fig 4). The results indicated that treatment with metformin could reverse the changes in adipogenesis seen in MGO / GAD treated cultures (Fig 5C).

The autophagy process was monitored by staining for LC3 II expression, a marker protein for auto-phagosome vesicles and provides the outcome as a puncta assay. This study presents the first observation of autophagy in carbonyl compound treated adipocytes. We analyzed the increase in puncta (auto - phagosomes) at the single-cell level, which were co-stained with GRP78. The results indicated that the number of autophagy puncta is greater in cells treated with MGO / GAD than the values in control cells, with increases from 8.32 ± 6.99 LC3 puncta/cell (N = 89) in the control to 12.38 ± 9.3 (N = 57) and 17.28 ± 13.11 (N = 68) after treatment with MGO and GAD respectively (Fig 5E). These results indicate that exposure to carbonyl compounds impairs the efficiency of the elimination pathways i.e., autophagy.

Interestingly, the addition of MET significantly decreased the number of LC3 puncta seen in the GAD treated cells by a factor of 3.7 and restored the values to control levels (MET+GAD: 4.69 ± 4.24, N = 85; GAD: 17.28±13.11, N= 68; control 8.32±6.99, N = 89). The results of cultures treated with only MET (6.45 ± 6.64 LC3 puncta/cell, N = 96) were similar to control group (Fig 5D-E). The immune staining and punctate measurements indicate that MET improves the autophagy function under glycated conditions. These results were verified by western blot (WB) analysis of the autophagy process efficiency based on LC3 I/II and p62. Fig 5F demonstrates an increase in P62 after treatment with the carbonyl compounds, while the addition of MET restored control levels of expression. These findings confirm the puncta analysis results and support the hypothesis that AGEs affect autophagy and impair the efficiency of the elimination pathways (Fig. 5E). Using spectroscopy, we were able to quantify the level of protein misfolding in cell lysates in the presence of ThT. Spectroscopy results indicate elevated levels of ThT fluorescence in the presence of MGO and GAD, which reflect increased amyloidogenic protein formation, correlated with impaired autophagy. The findings that in presence of carbonyl metabolites we monitored an elevation of amyloid levels by approximately 50% compared to the control, while metformin treatment restores control values (Fig 5G-H), demonstrate that hyperglycemic treatment reduces the efficiency of autophagy and elimination of amyloid aggregates while metformin restores normal values.

### Amyloid formation in adipose tissue

The protein misfolding associated with LDs was observed in cell cultures and was validated by multiple assays. We also demonstrated the appearance of amyloid proteins associated with LDs membrane in white adipose tissue (WAT; Fig 6B-C) and freshly isolated mature adipocytes (Fig 6E-F) by 4G8 antibody IF staining, or ThT staining visualized by confocal and polarized microscopies. To co-localize the potential alteration to a specific protein, we performed a WB with the 4G8 antibody on protein extracted from the WAT (Fig 6D) revealed a ≈55kDa protein and a high molecular weight >150 kDa which suggested for protein aggregates.

**Figure 6:**
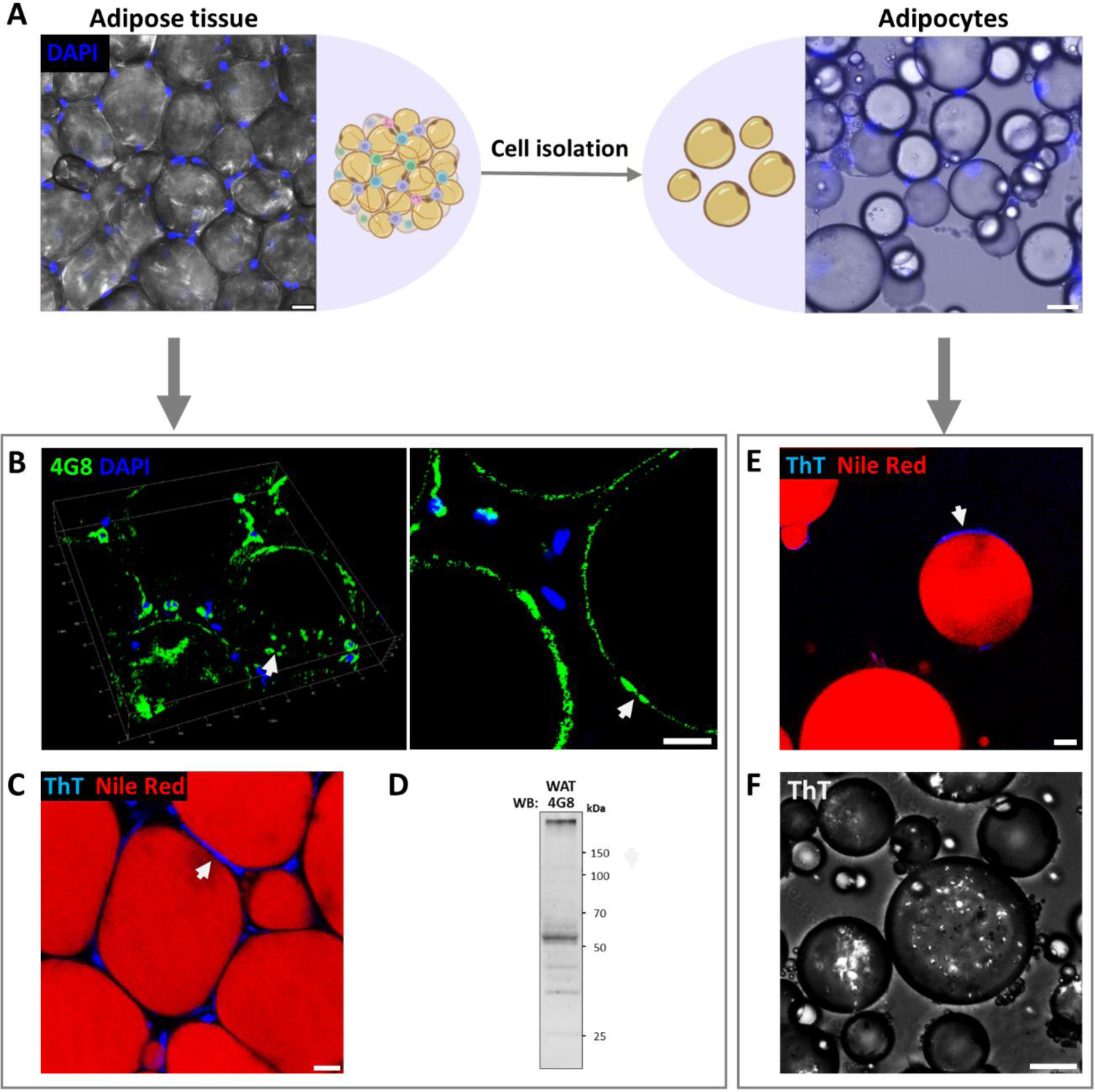
Lipid droplet associated protein aggregation in mouse adipose tissue and isolated adipocytes. **A.** Adipose tissue (WAT) and primary adipocytes visualized before and after cells isolation (Scale bar: adipose tissue = 20 µm, adipocytes = 50 µm). **B.** 3D confocal image (left image) of lipid droplet associated aggregates (white arrows), stained by 4G8 antibody. The arrows mark the same region on the LD surface in both images. (WAT; Magnification of X200; Scale bar = 15 µm)**. C.** Lipid droplet associated aggregates (white arrow) visualized by ThT staining (blue) and lipid droplets stained by Nile-Red. (WAT; Magnification of X200; Scale bar = 15 µm)**. D.** Immunoblotting of WAT lysate with 4G8 antibody. **E.** Fluorescence microscopy of isolated primary adipocytes stained with ThT (amyloids, white arrow) and Nile-Red (lipid content) (Magnification x200; Scale bar = 15 µm). **F.** Polarized light microscopy of ThT stained isolated adipocytes (Scale bar = 60 µm).

Therefore, bioinformatics tools were utilized to identify the LDs’ membrane proteins responsible for such alteration in protein structure. Protein alterations rely on their amino acid sequences as not all proteins become amyloidogenic. The candidate proteins related to LDs that potentially undergo misfolding were screened for aggregation-prone regions. AGGRESCAN tool allows to predict “hot-spot” areas along the protein sequence with the potential for amyloid aggregation. Comparison of the THSAr scores of LDs related proteins to known amyloidogenic proteins (Fig 7A), revealed that 6 out of the top 10 LDs proteins, namely ATGL, ABHD5, HSL, FSP27, DGAT2, and CIDE-A have a high amyloidogenic score (> 0.10). Taking this result together with the molecular size of the protein detected by the western blot (Fig 6D), it suggests that the observed protein is limited to a 55 kDa size, thus identifying the candidate protein as ATGL (Fig 7A). The hot spot profile of ATGL is presented in Fig 7B alongside amylin and APOE, which are known to form amyloid structures, with high score for amyloid formation in specific regions (Fig 7B). In relation to ATGL protein structure, comparison of 2D and 3D AGGRESCAN scores reveals a considerable overlap of the amyloidogenic regions predicted by these methods. 3D AGGRESCAN revealed several regions in the ATGL sequence that are prone to amyloid formation, namely around 140-160 and 330-360 AA have a particularly high correlation with the 2D and 3D prediction and are thought to affect the protein 3D structure (Fig 7C). Additionally, prediction tools PASTA-2, Fold Amyloid, MetAmyl, and WALTZ also confirm the identified sequence regions repeatedly emerge as hot-spots for amyloid formation (Fig 7D).

**Figure 7:**
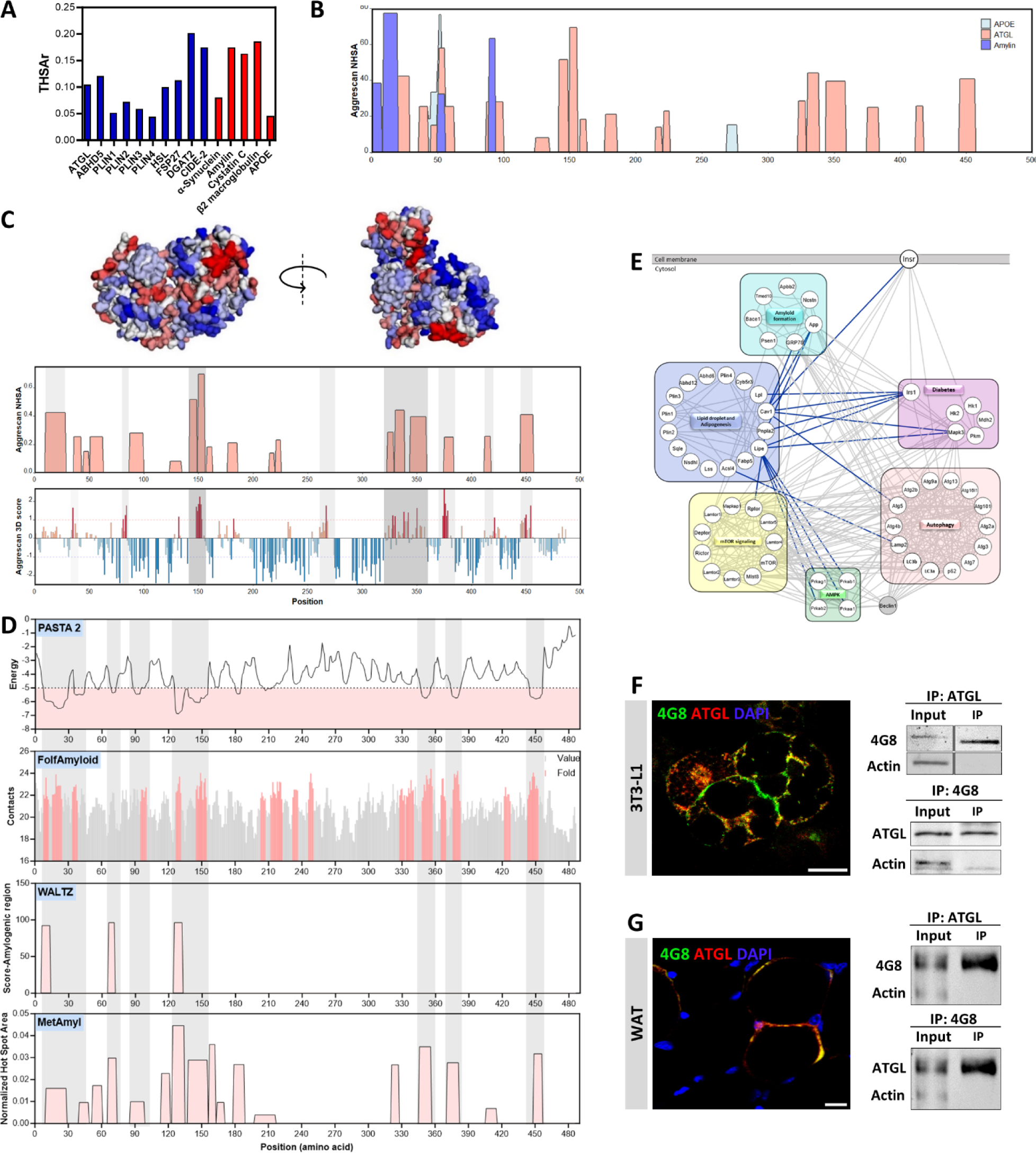
Prediction of lipid droplet associated protein that promotes amyloid secondary structure. **A.** AGGRESCAN THSAr score for protein amyloid aggregation of lipid droplet-associated (blue) compared with known amyloid forming proteins (red). **B.** The protein sequences of ATGL (red), APOE (light blue), and Amylin (blue) plotted by the AGGRESCAN aggregation “hot spots” tool. **C.** 3D AGGRESCAN score, alignment correlation of susceptible regions for amyloid formation in 2D and 3D schemes of ATGL protein. **D.** Predicted amyloid formation prone regions (red) of ATGL using PASTA 2 (energy<-5), FoldAmyloid, WALTZ and MetAmyl. Regions of the sequence that show correlation across three different algorithms are marked. **E.** STRING proteins association network categorized by cellular function and location. The connection between “lipid droplet and adipogenesis” to other categories are marked in blue. (Pnpla2 = ATGL). **F.** IF co-staining of ATGL and 4G8 antibodies in 3T3-L1 adipocytes (left; magnification of X630; Scale bar = 20 µm). Co-immunoprecipitation of ATGL and 4G8 and WB with 4G8 and ATGL respectively (right). **G.** IF staining in WAT for ATGL and 4G8 (left; magnification of X200; Scale bar = 15 µm). Co-immunoprecipitation of ATGL and 4G8 and WB with 4G8 or ATGL in WAT protein extraction (right).

In Fig 7E, STRING network analysis allows to demonstrate the linkage between amyloids, autophagy, and LD-related proteins. The presented proteins interaction network reveals that the proteins are clustered into four groups with associations with: lipid droplets, adenosine monophosphate-activated protein kinase (AMPK), autophagy, or mTOR signaling and there are functional connections between these groups. For example, the insulin receptor (Insr) signaling pathway (Irs1/IRS1-insulin receptor substrate) is connected to the autophagy through p62 (SQSTM1) that is an adaptor protein for selective autophagy of ubiquitinated protein aggregates, and is commonly used to assess the autophagy flux (Fig 5). P62 is also correlated with the degradation of amyloid aggregates because of the formation of misfolded proteins that p62 can recognize. In these networks, ATGL (Pnpla2), is a prominent protein of the lipid droplet cluster, and STRING analysis connects this protein to diabetes related pathways including mTOR signaling and amyloid formation.

The results of the bioinformatics (Fig 7) and western-blot analyses (Fig 6D) led us to focus on ATGL as the candidate protein for the amyloidogenic alterations observed in the LDs. The 4G8 antibody recognizes amyloid structures in other proteins and was used here to confirm the amyloid nature of ATGL, as demonstrated by IF and CO-IP of ALGL and 4G8 (Fig 7F, G). The co-staining of ATGL and 4G8 in cultured adipocytes (Fig 7F) and WAT (Fig 7G) result with co-localization (yellow area) of ATGL with 4G8, overlap regions suggest the ATGL conformational changes. The result was strengthened by Co-IP and WB assays with 4G8 antibody that was positive to ATGL and inversely, in both cell culture and WAT (Fig 7F-G). Our findings confirmed that ATGL is undergoing amyloid alterations in both 3T3-L1 adipocytes and WAT.

In this study, we present novel findings that shed light on the ATGL, a key protein in the formation of β-amyloid secondary structure that is associated with LDs. This discovery unveils a previously unknown link between ATGL and amyloidogenic protein changes in adipose tissue, offering valuable insight into potential mechanisms underlying diabetes-related complications and serve as a new platform that confirms the hypothesis of protein glycation in T2D.

## Discussion

Exposure to hyperglycemia leads to insulin resistance and metabolic impairment. Prolonged hyperglycemia results in the formation of glucose carbonyl metabolites and subsequently to the development of AGEs through protein glycation. Carbonyl metabolites, such as MGO and GAD, cause protein misfolding and structural changes that can affect protein functionality (14, 32, 33). Since adipocytes are central players in glucose metabolism, they were selected as a model system to study the effect of carbonyl compounds on protein alterations, and the deposition of β-amyloid structures formation. To this end, we examined the formation of amyloid formed on LDs in adipocytes cell cultures in the presence of MGO and GAD and WAT. We also examined the effect on autophagy in order to understand whether the observed accumulation of misfolded glycated proteins can be attributed to a failure of the physiological elimination mechanisms.

Metformin (MET), prescribed as a first-line treatment for T2D, has also been shown to decrease serum levels of MGO by about 30% in humans (34). MET is thought to influence the MGO level through a direct chemical interaction between the two molecules (MGO-MET), or MET effect on glycemic control that inhibit MGO production (35, 36). The effect of MET on GAD treated cells has so far been reported only in-vitro study on macrophage cell line. In other aspect, the effect of GAD on LDL (low-density lipoprotein) glycation leads to cholesterol accumulation in macrophages and this process was inhibited by MET (37). Previous studies describe the effect of MET on adipocyte differentiation, and regulation of ER-stress and AMPK signaling (38–40). The results present the beneficial impact of MET on autophagy in MGO and GAD treated adipocytes, autophagy affected by AMPK signaling (Fig 5). In addition, MET restores the level of adipogenesis after this has been reduced by exposure to MGO and GAD and lowering cellular amyloid level that were elevated by the carbonyl compounds (Fig 5G). In this study, profound effects were observed using low concentrations of MET, whereas other research involving 3T3-L1 cells required higher doses to impact adipogenesis and AMPK signaling pathways (38, 39). As suggested in Fig 7E, the STRING network connected the AMPK role on diverse cellular processes including autophagy, mTOR signaling, as well as a correlation with autophagy and lipolysis. mTOR is connected to the hexokinase 2 enzyme (Hk2), which phosphorylates glucose to produce glucose-6-phosphate for glycolysis and is upregulated in T2D.

Our data reveal the effect of AGEs formed in 3T3-L1 cells on molecular vibrations, as observed in the RAMAN spectra peaks that reflect secondary structures. The RAMAN spectroscopy is a sensitive technique that provides a biochemical “fingerprint” to resolve the structural molecular changes in the intracellular compartment (41, 42). Here, we identify the changes in the intensity of the amide I and III peaks as a response to hyperglycemia in adipocytes. These changes reflect alterations in protein conformation and an increase in β-sheet structures (Fig 4). Such changes in the 1655 and 1260 cm-1 peaks were previously described in primary mice neurons after exposure to amyloid-β protein oligomers (43), and β-sheet formed protein accumulation in pancreatic cancer cells (44). The RAMAN results are well correlated with the ThT spectroscopy (Fig 5G), staining and polarized microscopy (Fig 3) that emphasize the LD-associated amyloid aggregations.

Lipid droplet-associated proteins are a group of proteins that have been shown to be critical in regulating lipid metabolism. Among this group, ATGL has emerged as a central player in the regulation of lipolysis and triglyceride (TG) metabolism. ATGL has been shown to be essential for efficient lipid mobilization from lipid droplets, particularly in response to energy demands (45, 46). In addition, recent studies have demonstrated that ATGL may also be involved in the regulation of autophagy-mediated lipid degradation by the LC3-interacting region (LIR) motif (47). Our results demonstrate the negative effect of AGEs on both lipolysis and autophagy (Figs 2 and 5). Interactions between LDs associated proteins and autophagy - related proteins suggest that these proteins play a critical role in maintaining cellular lipid homeostasis. ATGL knockout mice, which lack this key player in the lipolysis process, tend to exhibit moderate obesity (48, 49). This suggested to be correlated with the impaired lipolysis observed in adipocytes following treatment with MGO and GAD, accompanied by an elevation in the formation of amyloid structures associated with lipid droplets (Figs 2, 3). Our findings underline the significant role of ATGL in the lipolysis process therefore suggest the disruption of its function could be intensified by the formation of β-amyloid structures on proteins associated with lipid droplets.

Amyloid-forming proteins, like amylin in the pancreas, play a role in the development of type 2 diabetes (T2D) when they form amyloid fibrils (50–52). These amyloid deposits accumulate in the pancreas, thereby impairing β cell function and disrupting glucose homeostasis. Interestingly, there is a high comorbidity between diabetes and other amyloid-related diseases, such as Alzheimer’s disease and systemic amyloidosis (53–55). This suggests the presence of common mechanisms and processes involving amyloid formation. Understanding these mechanisms is crucial for developing treatments for amyloid-related diseases. In the context of T2D and amyloidosis, we propose to explore adipocytes as a novel focal point for investigation.

In summary, the results of the current study highlight the detrimental effects of hyperglycemia-induced glucose carbonyl metabolites and AGEs formation on adipocyte differentiation and metabolism. Such condition promotes protein misfolding and the formation of β-amyloid deposits. Furthermore, our results suggest that targeting lipid droplet-associated proteins, such as ATGL, may hold promise for developing therapeutic interventions for metabolic disorders. The presented findings could potentially serve as valuable markers for gaining deeper insights into metabolic changes occurring in adipose tissue and is a significant step forward in understanding the role of ATGL in cellular dysfunction, investigating the interplay between AGEs formation, amyloidosis and T2D. Further research in this area is necessary to unravel the underlying mechanisms and explore potential treatment strategies for amyloid-related diseases.

## Materials and Methods

### Visceral adipose tissues (VAT)

Epididymal white visceral adipose tissue (WAT) were collected from C57BL/6J mice (Envigo RMS Limited, Jerusalem, Israel) and immediately used as fresh, frozen (by liquid nitrogen) or for mature adipocyte isolation following collagenase digestion as previously described (56, 57). The mice were kept in a conventional facility of Tel Aviv University (TAU) with 12 h light/dark cycles and were fed with standard chow diet and water were provided ad libitum. Animal care and experiments were in accordance with the guidelines of the IACUC Approval (01-21-044). **Mature adipocytes:** Were isolated from VAT as described (56). Briefly, the tissue was ground to a fine consistency in HBSS solution (Biological Industries, Israel), then incubated with collagenase solution (Sigma-Aldrich, c-5138) for one hour at 37^◦^C with shaking and the digested tissue was filtered through a 100 µm cell strainer (SPL life s cience) and centrifuged at 1800 rpm for 5 min. The mature adipocytes were collected for ThT staining as explained below.

### Cell culture, adipogenic differentiation and hyperglycemia cell growth

3T3L1 cells (ATCC, USA) were seeded at 1×10^4^ cells/cm2 in a growth medium (GM, Dulbecco’s modified Eagle’s medium DMEM, 4.5 mg/ml glucose; Gibco™) with 10% fetal bovine serum and 1% L-glutamine (Biological Industries), 0.1% penicillin–streptomycin (P3032, 85555; Sigma-Aldrich), 0.5% 4-(2-hydroxyethyl)-1-piperazinee thane-sulfonic acid (HEPES; Biological Industries). Differentiation medium (DM) supplemented with 100 IU/mL insulin (41-975-100; Biological Industries, Israel), 1μM dexamethasone (Sigma-Aldrich), and 400μM 3-isobutyl-1-methylxanthine (IBMX; 2885842; BioGems) for two days then replaced by a supporting medium (SM); consisting of the GM supplemented with 100 IU/mL insulin (58, 59). Once adipogenesis was visualized in the cells, carbonyl compounds - methylglyoxal (MGO, M0252) or glycolaldehyde (GAD, G6805) both from Sigma-Aldrich were added to the medium in a final concentration of 0.001 mg/ml or 0.01mg/ml, respectively for 14 days and in some cultures 5µM metformin (MET) (BioVision, 1691) added for a period of 6 days. The cultures incubated at 37°C in humidified atmosphere containing 5% CO2 .

### Morphological features to follow differentiation

Cells growth and differentiation process was followed by morphology changes, LDs accumulation monitored on live cultures. The adipogenesis quantification at the macro-scale X40 for mapping the level of adipogenesis (LOA) follow up using phase-contrast EVOS FL Auto-2 Microscope (Invitrogen) (60). Images at higher magnification x200 allow to analyze the cell and LDs size (59).

### Bioinformatics

The functional connection between proteins analyzed by STRING software https://string-db.org/ site (61). Cytoscape software platform used for visualizing the STRING networks (62).

### In silico aggregation analysis

AGGRESCAN server (http://bioinf.uab.es/aggrescan/) (63) was employed to predict hot-spot aggregation regions in various LD-bound proteins. The server utilizes an algorithm that assesses the potential for amyloid aggregation based on the input protein sequence. Their THSAr (Total hot spot area per residue) score was calculated to quantitatively evaluate the amyloidogenic potential of the target proteins. This score provides an indication of the protein’s propensity for amyloid aggregation. AGGRESCAN 3D server was used to predict potential aggregation regions of ATGL (http://biocomp.chem.uw.edu.pl/A3D2/) (64). This tool allows the visualization of predicted hot-spot aggregation regions in three dimensions. Complementary aggregation property predictions were also performed using the WALTZ (https://waltz.switchlab.org/index.cgi), PASTA2 (65), MetAmyl (66), and FoldAmyloid (67) algorithms that focus on amyloidogenic regions predictions of ATGL sequence (Pnpla2 protein [Mus musculus]; Accession: UniProt Q8BJ56-1).

### Cell morphology, staining and imaging

#### 1. Whole-Mount immunofluorescence staining

VAT whole-mount staining was performed as previously described (58). Primary antibodies: ATGL (mouse, Santa Cruz sc-365278), 4G8 (Beta Amyloid 17-24; mouse, Bio Legend, SIG-39240). Secondary antibodies: Alexa 488 (A 21141), Alexa 555 (A-21127) from Invitrogen, mounting medium Fluoroshield™ containing 4’, 6-diamidino-2-phenylindole (DAPI) (Electron Microscopy Sciences, #17985-10) and viewed by confocal microscope (Leica SP8; Leica, Wetzlar, Germany).

#### 2. Immuno-fluorescence (IF) cell staining

Cells were fixed using 4% paraformaldehyde containing 0.03M sucrose in phosphate-buffered saline (PBS) for 10 minutes at RT, then washed in 1% PBS with 0.5% triton. Blocking solution, Tris buffer saline (TBS) 1% containing 1% normal goat serum and 1% bovine serum albumin (BSA). The cells incubated with primary antibodies: LC3b (rabbit, Abcam ab192890; mouse, Sc-271625), GRP78 (rat; SC-13539), perilipin 1 (mouse; SC-390169), GLUT4 (mouse; SC-53566), ATGL (mouse; Santa Cruz SC-365278), 4G8 (Beta Amyloid 17-24; mouse, Bio Legend, SIG-39240). Secondary antibodies; Anti-mouse: Alexa 488 (A21121 and A 21141), Alexa 555 (A-21127) from Invitrogen, Anti-rabbit: Cy3 (Jackson Immuno Research Laboratories, Inc.), Anti-rat: donkey anti-rat Alexa 647 (Invitrogen). The stained cells were mounted with fluoro-Gel mounting medium containing 4’, 6-diamidino-2-phenylindole (DAPI) (17985-50, Electron Microscopy Sciences). Cells visualized and photographed by fluorescence microscopy (Nikon, Eclipse Ci) or by Zeiss LSM-710 confocal microscope and analyzed using ImageJ software (NIH, Bethesda, MD, USA).

#### 3. Glucose uptake assay

using a fluorescent glucose analog, 2-NBDG (2-Deoxy-2-[(7-nitro-2, 1, 3-benzoxadiazol-4-yl) amino]-D-glucose) (11046, Cayman chemical). Cultured 3T3-L1 cells “starved” in a glucose-free medium at 37°C for 1 hr, then added 100µM 2-NBDG and insulin for 1 hr, washed with PBS and fresh media was added, cultures observed by live imaging (Incucyte® SX5, Sartorius).

#### 4. Fluorescent staining for amyloids and lipid droplets

Fixed adipose tissue, isolated adipocytes, or cultured adipocytes were incubated with 5 mM Thioflavin T (ThT; Sigma T3516), 7mM Congo red (CR; T6277, Sigma) and 10µg/ml Nile Red (Sigma N-3013) for 30 min in room temperature, then were mounted with fluoroshield mounting Fluoroshield™ (Electron Microscopy Sciences, #17985-10, without DAPI), images were acquired by confocal microscope (Leica SP8; Leica, Wetzlar, Germany). ThT and CR-stained cells observed, and images acquired using polarized light microscope (Nikon, Japan), fluorescence microscope (Eclipse Ci; Nikon), EVOS FL Auto-2 Microscope (Invitrogen), confocal microscope-Zeiss LSM-710, Chameleon 720 (690-1064) laser and images were analyzed by ImageJ.

#### 5. Transmission Electron Microscopy (TEM)

Cells were fixed overnight in 2.5% Glutaraldehyde in phosphate-buffered (PBS) at 4 °C was then washed several times with PBS and post-fixed in 1% OsO4 in PBS for 2 h at 4 °C. Dehydration in graded ethanol and embedded in Glycid ether for preparation of thin sections mounted on Formvar/Carbon coated grids. Sections were stained with uranyl acetate and lead citrate and examined using a JEM 1400 Plus transmission electron microscope (Joel, Japan). Images were captured using SIS Mega view III and the TEM imaging platform (Olympus).

#### 6. RAMAN microscopy and spectroscopy

Adipocytes were treated with MGO and GAD for 11 days and then cells were re-plated on 60nm gold-coated glass coverslips for two days. For the RAMAN spectroscopy, cultures were washed with PBS and air-dried. Cells were excited with a 532 nm laser and the spectral range was 1000-1800 cm-1, with maximal laser power of 10mW and exposure time of 60 seconds collected the RAMAN spectral recorded by Lab Ram HR of Horiba Jobin Yvon (NewRoad, Olympus, Japan). Images were acquired by objective x 50 (N.A=0.55), and the spectra were collected by objective x100 (N.A=0.9) (Olympus, Japan). The RAMAN signal collected by the CCD detector cooled down to -70° and RAMAN spectra processing was obtained using SpyctraGryph software .

#### 7. ThT spectroscopy

Cells harvested from cultures were lysate in 25mM Tris pH 7.4, 150mM NaCl buffer containing 0.5% sodium deoxycholate, 0.1% SDS, and protease inhibitors: (phenylmethylsulfonyl fluoride, PMSF 1mM; 1-chloro-3-tosylamido-4-phenyl-2-butanone, TPCK, 10 μg/ml; aprotinin, 10 μg/ml Sigma) incubated for 30min on ice. Thioflavin T (ThT; T3516, Sigma-Aldrich) 30 µM was added to samples in 96-well black microplate measured at excitation 420 nm/emission 485 nm using a Synergy™ H1 multi-mode microplate reader (BioTek Instruments, USA) (14).

#### 8. Lipolysis

3T3-L1 adipocytes cultured and treated with MGO and GAD as described. For the lipolysis induction, 10 μM of Forskolin (PeproTech) was added to the supporting medium; the forskolin was kept for 3 hr. as described by us (30). Live imaging and visualization were performed using EVOS FL Auto-2 Microscope (Invitrogen).

### Biochemistry and protein identification by western blot (WB)

Protein extraction or immunoprecipitation (IP) using Protein A/G PLUS-Agarose (Santa Cruz biotechnology; sc-2003) (58). The protein samples were re-suspended in Lamelli buffer and separated on SDS polyacrylamide gel electrophoresis gel, then transferred to nitrocellulose. Nitrocellulose membranes incubated with primary antibodies against LC3b (rabbit, Abcam ab-192890), p62 (rabbit, Abcam ab-109012) ATGL (mouse, Santa Cruz sc-365278), 4G8 (mouse, Bio Legend, SIG-39240), and actin (mouse, MP-691001/2). Secondary antibodies horseradish peroxidase (HRP) conjugated used; bovine anti-rabbit (SantaCruze; sc-2370), goat anti-mouse (Jackson; 115035). Signal detected with chemiluminescent substrate (Super Signal ^TM^ West Pico PLUS, 34578, Thermo Fisher) exposed and quantified digitally on Fusion FX7 (Vilber Lourmat) .

### Gene expression (qPCR)

As previously described (25), total RNA was extracted from 3T3-L1 cells using Trizol reagent (Bio Tri RNA; Bio-Lab Ltd., Jerusalem, Israel) and reverse transcribed to cDNA using a high-capacity cDNA reverse transcription kit (Applied Biosystem, Waltham, MA, USA). Transcript levels were measured with SYBR green (Applied Biosystem, Waltham, MA, USA) using STEPONE plus system (Thermo Fisher Scientific, Waltham, MA, USA). All data were normalized to actin by the delta-delta Ct method (68). For qPCR amplification, we used the primers for the following genes:

> Actin F-CATCGTGGGCCGCCCTAGGCACCA; R-CGGTTGGCCTTAGGGTTCAGGGGG
>
> Glut4 F-TTCACGTTGGTCTCGGTGCT; R-TAGCTCATGGCTGGAACCCG.

### Statistical analysis

Statistical analysis was performed using GraphPad Prism 9 software (La Jolla, CA). One-way ANOVA or two-way ANOVA tests with Turkey post hoc test were performed in accordance with variable number, p-value <0.05 was considered as statistically significant. All error bars shown in the figures are standard deviation (SD). Data plotted on GraphPad Prism 9 software and Microsoft Office Excel-2013.

### Schematic illustrations

were created by the Bio Render software https://biorender.com.

## Acknowledgments

Roza Izgilov was partially supported by a Scholarship from the Tel Aviv University Healthy Longevity Research Center. Part of this work was performed in fulfillment of the requirements for a Ph.D. degree of Roza Izgilov, Faculty of medicine, Tel Aviv University, Israel. Adam Noam Gophna, a student from the Alpha Research Program, for imaging and analysis part of Fig 3. Shira Eidelheit for analysis part of Fig 2. Ann Avron for the editorial assistant.

